# A modular HaloTag platform for engineering magnetically responsive bacterial microrobots

**DOI:** 10.64898/2026.01.25.701580

**Authors:** Xiang Wang, Pascal Poc, Elena Totter, Simone Schuerle

## Abstract

Functionalizing bacteria with magnetic nanoparticles (MNPs) is a common route toward magnetically responsive bacterial microrobots. However, existing strategies are often limited by low functionalization efficiency, weak binding, poor reproducibility, and nonspecific interactions. Here, we present a robust, specific, and reproducible magnetic functionalization platform based on the self-labeling protein HaloTag, yielding magnetic bacterial microrobots termed HaloBots. Using *Escherichia coli* as an engineering host, HaloTag was displayed on the outer membrane via the Lpp–OmpA anchoring system, with the assistance of a Long and Flexible (LF) peptide linker. Optimization of genetic engineering vectors and colony purification enabled robust and uniform HaloTag display across bacterial populations. In addition, MNPs were modified with chloroalkane–PEG ligands to enable site-specific covalent conjugation with HaloTag. Tuning the ligand density on MNPs revealed a critical balance between ligand accessibility and surface charge for achieving efficient and specific attachment. Consequently, the resulting HaloBots exhibited stable MNP conjugation and reliable magnetic actuation. Collectively, this work establishes a modular and tunable strategy for engineering magnetically responsive bacterial microrobots.

## Introduction

Magnetic microrobots have gained significant attention due to their potential for precise remote control under magnetic fields.^1-4^ Notably, biohybrid magnetic microrobots are an important subclass that uses microorganisms as controllable living biological components, providing on board propulsion and sensing.^5-7^ These living microrobots can be guided by external magnetic fields and have been explored for diverse applications, including cancer therapy,^5,8,9^ targeted drug delivery,^7,10^ antiviral therapy,^11^ and biosensing and diagnostics.^12,13^ Bacteria are a frequently used class among the utilized microorganisms, and among which magnetotactic bacteria (MTB) have long served as model hosts for magnetic biohybrid microrobots. They inherently exhibit a strong and stable magnetic response due to their ability to naturally biomineralize magnetosomes, which are chains of iron oxide nanoparticles wrapped in a lipid bilayer.^14^ In contrast, more commonly used yet non-magnetic bacterial strains possess advantages such as metabolic versatility, and genetic tractability which render them particularly attractive candidates for the development of biohybrid magnetic microrobots once magnetically functionalized. To render non-magnetic bacteria magnetic, akin to MTB, genetic modification strategies, such as engineered ferritin expression,^15,16^ magnetosome gene transfer,^17,18^, have been explored. However, these approaches have so far yielded only limited magnetic responses and impose a substantial genetic burden on the host. As a result, current strategies mainly focus on conjugating magnetic nanoparticles (MNPs) to the bacterial surface. Established approaches include electrostatic adsorption,^19,20^ Biotin-Streptavidin complexation,^7^ and bioorthogonal click chemistry.^21^ While these methods have demonstrated MNP attachment on bacterial surface and proof-of-concept magnetic actuation, several major challenges hinder their translation from bench to bedside. From an engineering perspective, bacterial microrobots designed for translational applications should meet several key criteria. First, the conjugated MNPs must be present in sufficient numbers to enable reliable magnetic actuation at both the single-cell and swarm levels. Furthermore, attachment methods should be highly specific to avoid nonspecific binding and off-target interactions. Finally, the conjugation must be stable under physiological conditions to ensure sustained functionality in vivo. Electrostatic adsorption allows for high MNP loading but is inherently nonspecific, and electrostatic forces are often unstable and can be affected by various factors such as temperature and ionic strengths.^22-24^ Biotin–streptavidin complexation provides high specificity, yet in physiological environments it may still lead to nonspecific interactions,^25^ and achieving homogeneous and reproducible nano–bio conjugation remains challenging.^26,27^ Bioorthogonal click chemistry provides strong covalent coupling, yet often requires prior modification of the bacterial surface,^28,29^ and our recent study has demonstrated the need of pretreatment of bacterial surface to achieve high click reaction efficiency.^30^ Therefore, developing new strategies that combine high loading efficiency, specificity, stability, and simplicity will be essential for advancing bacterial magnetic microrobots toward practical applications.

Self-labeling proteins such as HaloTag,^31,32^ SNAP-tag,^33,34^ and CLIP-tag^35^ provide powerful tools for labeling and functionalizing living cells. They have been widely used for applications including protein imaging^36^, protein–protein interaction studies^37^, and targeted delivery of synthetic molecules^38^. Among them, HaloTag, an engineered haloalkane dehalogenase, has become particularly attractive because it can covalently react with ligands containing chloroalkane moiety with fast labeling kinetics.^39,40^ This reaction is not only efficient, but also stable due to formation of a covalent bond, simple to perform, and robust under physiological conditions, making HaloTag especially convenient over all the self-labeling tags. In addition, chloroalkane-modified ligands are readily accessible, either through commercial sources or straightforward synthesis, providing a versatile plug-and-play platform for covalent attachment of diverse cargos, including fluorescent dyes,^41^ biomacromolecules,^42^ polymers,^43^ and nanoparticles.^44^ When genetically expressed and displayed on the bacterial surface, HaloTag can thus serve as a universal docking module that provides stable and reproducible functionalization while preserving cell viability. Although HaloTag surface display has not yet been systematically demonstrated across different bacterial species, established protein display systems such as the Lipoprotein–Outer membrane protein (Lpp–OmpA) in Gram-negative bacteria provide a practical way to present HaloTag on the bacterial surface.^45^ By functionalizing MNPs with chloroalkane ligands and conjugating them to surface-displayed HaloTag, this approach offers a promising strategy for engineering magnetic bacterial microrobots.

In this study, we introduce a HaloTag-mediated strategy for the magnetic functionalization of bacterial microrobots (**Fig. 1**). By genetically engineering the Gram-negative bacterium *Escherichia coli*, HaloTag proteins were efficiently displayed on the outer membrane, enabling site-specific and covalent conjugation with chloroalkane-modified magnetic nanoparticles (CA-MNPs). This approach allows MNPs to be directly anchored to living bacteria without the need for additional surface treatment. The resulting functionalized bacteria, termed HaloBots, exhibit stable magnetic coupling and reliable torque-driven actuation under rotating magnetic fields. By combining genetic surface display with controlled chemical conjugation, this work establishes a robust and reproducible platform for the magnetic functionalization of bacterial microrobots.

**Figure 1.**
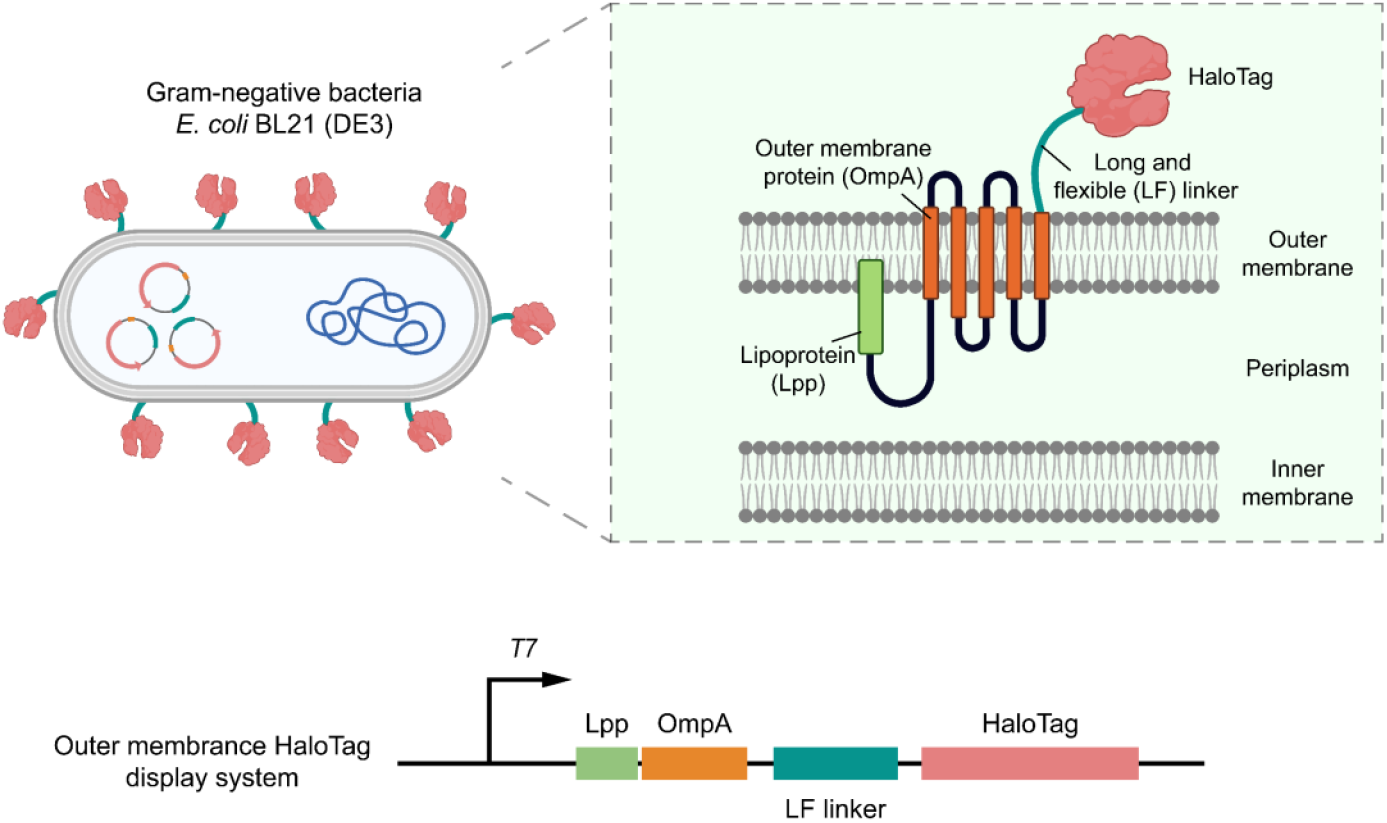
Outer membrane display system of HaloTag in Gram-negative bacteria. The system consists of several functional modules: The Lipoprotein (Lpp) acts as a signal peptide for secretion of the fused protein. The Outer membrane (OmpA) anchors to the membrane and extends passenger proteins to the cell surface. The Long and Flexible linker (LF linker) provides spatial separation between the outer membrane and the displayed HaloTag11 protein. The fusion construct was expressed using the engineered plasmid pWX118. Illustrations were generated with BioRender.com.

## Results

### Optimizing plasmid designs for surface display of HaloTag on *E. coli*

To begin our study, we first selected a suitable bacterial strain for HaloTag surface display. *E. coli* BL21 (DE3), one of the most widely used strains for expressing fusion proteins, was chosen as the engineering host. Gram-negative bacteria possess both an inner and an outer membrane, and for our purposes HaloTag needed to be displayed on the outer membrane. Since the molecular weight of HaloTag is around 33 kDa and it’s structurally simple, we employed the well-established and reliable lipoprotein (Lpp)–outer membrane protein A (OmpA) display system (**Fig. 1**), which has been reported to display similar size or larger proteins.^46,47^

To visualize the displayed HaloTag and evaluate the plasmid designs, HaloTag was labeled with a membrane-impermeable sulfo-Cyanine5 dye containing a chloroalkane (CA) group (CA-sCy5, **Fig. 2A**). CA-sCy5 was synthesized via an amide coupling reaction, in which CA-NHS was covalently linked to sulfo-Cy5-amine in the presence of N,N-Diisopropylethylamine (DIPEA) (Fig. S1-S3). For the initial attempt, we constructed plasmid pWX100 based on the constitutive vector pM965, which carries the *rpsM* promoter and GFP (**Fig. 2B**). Constitutive expression was chosen to simplify the workflow, avoiding the need for growth monitoring or inducer addition. In this construct, the GFP gene was replaced with HaloTag, positioned downstream of Lpp–OmpA to generate the Lpp–OmpA–HaloTag fusion. After transformation into *E. coli* BL21 (DE3) and subsequent labeling with the membrane-impermeable ligand, we imaged the bacteria and quantified the fluorescence intensity (**Fig. 2C and 2D**). However, the results indicated the amount of surface-displayed HaloTag was modest, suggesting that further optimization was required.

**Figure 2.**
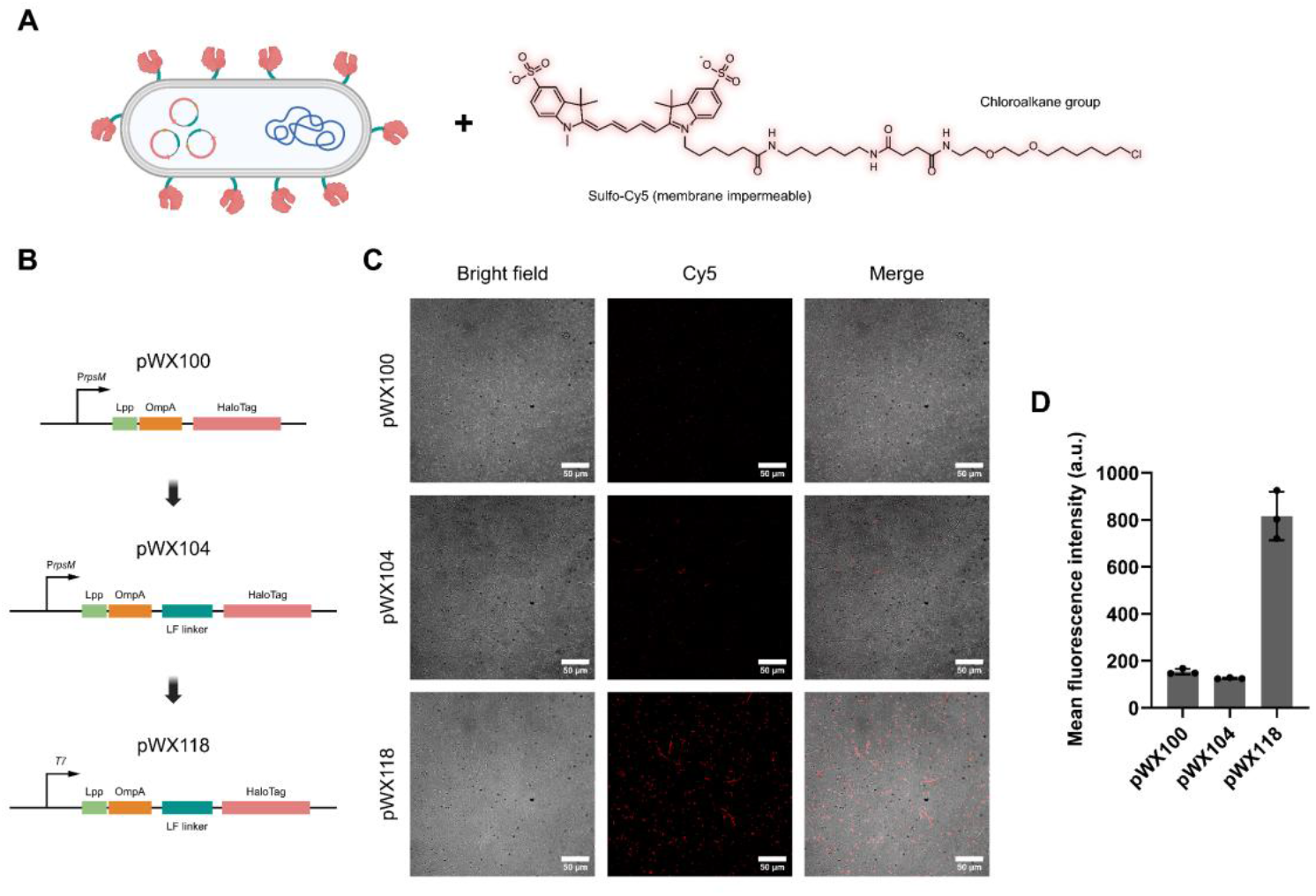
Optimizing plasmid designs for surface display of HaloTag on *E. coli*. **(A)** Schematic illustration of labeling process of surface-displayed HaloTag using a fluorescent ligand. The bacteria were incubated with Sulfo-Cy5 conjugated to a chloroalkane (CA) group, a HaloTag-specific and membrane-impermeable ligand, enabling selective labeling of surface-displayed HaloTag proteins. **(B)** Designs of plasmid constructs pWX100, pWX104, and pWX118. **(C)** Microscopy images of bacteria expressing HaloTag and labeled with fluorescent ligand. **(D)** Quantification of bacterial mean fluorescence intensity from microscopy images. Data are presented as means ± SD, *n* = 3 biological replicates.

Based on the design of the first construct, we hypothesized that steric hindrance might account for modest display efficiency. Proteins displayed on the outer membrane can be partially blocked by surrounding components such as lipopolysaccharides (LPS). To address this, we inserted a Long and Flexible (LF) peptide linker between OmpA and HaloTag (**Fig. 2B**). This linker was designed by combining a previously reported spacer peptide^48^ with glycine-rich repeats (GGGGS). Interestingly, with this modification, the resulting plasmid, pWX104, led to noticeably elongated bacterial cells (**Fig. 2C**). When exposed to elevated temperature, antimicrobial chemicals, and metabolic stress, bacteria often experience filamentous morphological changes.^49^ In our case, the elongation of the bacteria was likely caused by membrane stress due to overexpression of the large fusion protein, with the insertion of the LF linker. However, despite the morphological change, the HaloTag display efficiency of pWX104 remained similarly low compared to pWX100 (**Fig. 2D**).

We then realized that the constitutive expression system might not be robust enough to achieve high levels of HaloTag display on the bacterial surface. To further enhance expression, we switched to the pCDFDuet-1 vector, which carries a Lac-operon–inducible T7 promoter. The Lpp–OmpA–LF linker–HaloTag construct was cloned into the first multiple cloning site of this vector, generating plasmid pWX118 (**Fig. 2B**). After transformation into *E. coli*, expression of the fusion protein was induced with isopropyl β-d-1-thiogalactopyranoside (IPTG). As a result, plasmid pWX118 achieved substantially higher levels of HaloTag display compared to the previous designs (**Fig. 2C and 2D**). This improvement was not only due to the stronger inducible promoter but also to the choice of antibiotics. The earlier plasmids relied on ampicillin for selection, which is known to be relatively weak and less stable, whereas pWX118 was maintained under streptomycin selection, providing stronger and more consistent pressure. For these reasons, we selected plasmid pWX118 for the subsequent experiments.

### Colony purification improves efficiency of HaloTag surface display

Although plasmid pWX118 enabled high levels of HaloTag surface display, we noticed that the proportion of HaloTag-positive bacteria within the population was still limited, as indicated by the large number of non-labeled cells in **Fig. 2C**. After DNA transformation, a single colony consists of bacteria carrying different plasmid copy numbers or even cells that have lost the plasmid. In conventional bacterial engineering for soluble protein production, researchers primarily focus on total yield and therefore rarely examine the proportion of protein-expressing cells. In contrast, our goal requires a high fraction of bacteria displaying HaloTag on the membrane, as the functionality of the microrobots depends on the proportion of modified cells rather than on bulk protein production.

Therefore, after transformation, we re-streaked the colonies on LB–agar plates supplemented with streptomycin for another two times, to purify the bacterial clones and to achieve both a high and homogeneous plasmid copy number. The results showed that, using purified colonies and following induction of protein expression, nearly all bacteria displayed HaloTag on their surface (**Fig. 3A**), in clear contrast to the non-transformed bacteria (**Fig. 3B**). High-magnification imaging further confirmed that HaloTag proteins were localized on the surface of the bacteria (**Fig. 3C**). This purification step proved essential for achieving a homogeneous population, which is critical for the subsequent magnetic functionalization.

**Figure 3.**
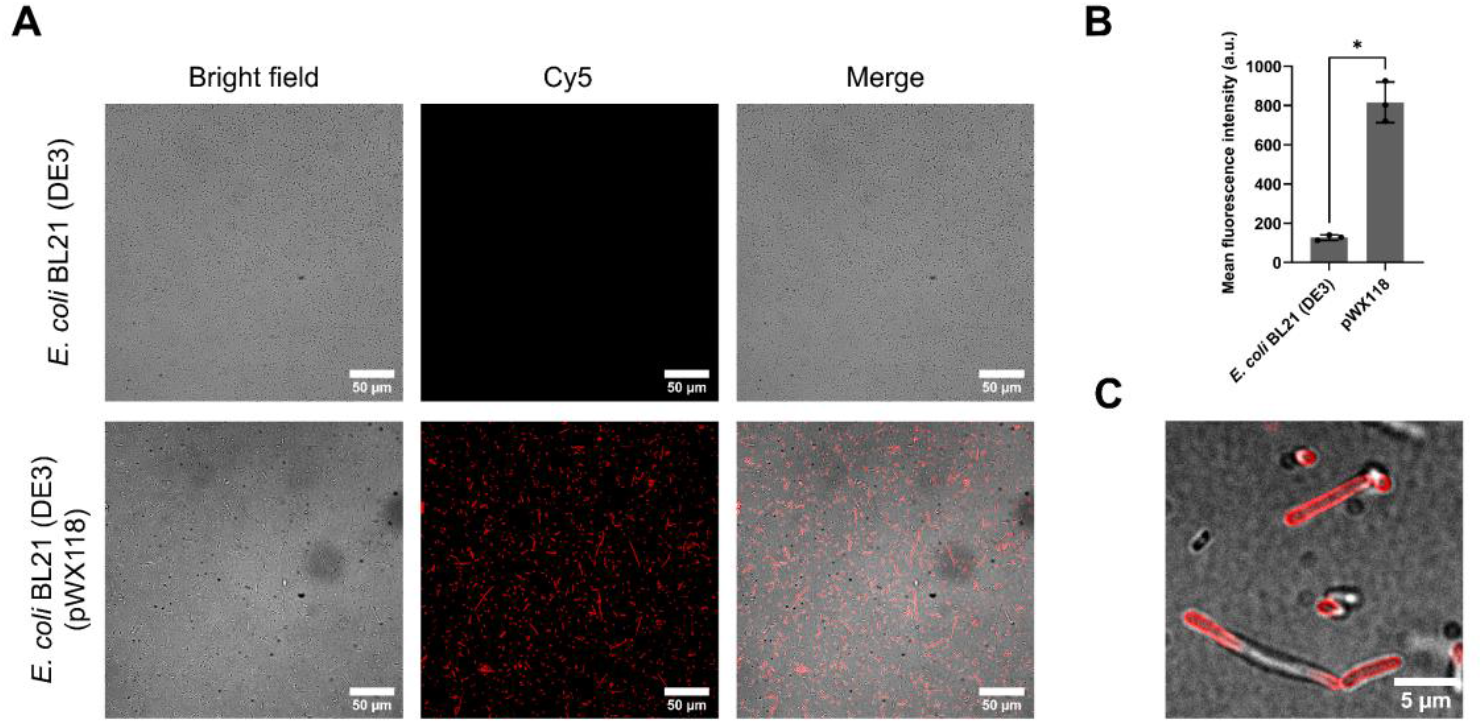
Fluorescence labeling of *E. coli* after colony purification. **(A)** Microscopy images of non-transformed bacteria and HaloTag-expressing bacteria labeled with the fluorescent HaloTag ligand. **(B)** Quantification of bacterial mean fluorescence intensity from microscopy images. Data are presented as means ± SD, *n* = 3 biological replicates. Statistical significance was determined using the nonparametric Mann-Whitney *U* test. \**P* ≤ 0.05. **(C)** High-magnification images showing surface localization of HaloTag on the bacterial membrane.

### Modification of magnetic nanoparticles (MNPs) with HaloTag ligand

Having established a robust HaloTag surface display, we next sought to prepare HaloTag-binding MNPs for bacterial magnetic functionalization. Similar to the fluorescent ligands, MNPs also needed to be modified with chloroalkane (CA) groups. In our preliminary studies, we functionalized MNPs with the same ligands previously used for fluorescent labeling, in which the sulfo-Cy5 dye was conjugated. To evaluate these particles, we tested them with *E. coli* (pWX118) displaying HaloTag. However, no functionalization could be observed, either under rotating magnetic fields (RMF) or by direct inspection of the bacterial pellets (data not shown). Considering that spatial accessibility is critical for the reaction between HaloTag and its ligands, and that MNPs impose considerably greater steric hindrance compared to small-molecule dyes, we introduced a long polyethylene glycol (PEG) linker (∼2000 Da) between the chloroalkane group and the MNP surface.

Prior to MNP functionalization, commercially available CA-COOH was activated to CA-NHS using TSTU and DIPEA, and subsequently conjugated to NH2-PEG_2000_-COOH to generate CA-PEG_2000_-COOH (Fig. S4-S9). The MNP modification was then initiated by activating this CA-PEG2000-COOH intermediate to its NHS-ester form, followed by coupling to the surface amine groups of commercial MNPs (**Fig. 4A**). In our initial attempts, we used a large excess of CA-PEG_2000_-NHS to modify amine-MNPs, in order to cover as many amine groups as possible. With these CA-modified MNPs, we occasionally observed positive functionalization results from TEM imaging (Fig. S10). However, the functionalization efficiency using these MNPs was low, as most bacteria were either not attached or carried only a few MNPs. Therefore we hypothesized that complete coverage of surface amine groups on MNPs by ligands may not be necessary. More importantly, previous studies have shown that excessive PEGylation of nanoparticles may create steric hindrance that blocks the access of biomolecules.^50^ Besides, a thick PEG layer can also mask the reactive CA groups, thereby reducing effective binding efficiency. Therefore, we sought to refine our MNP modification process.

**Figure 4.**
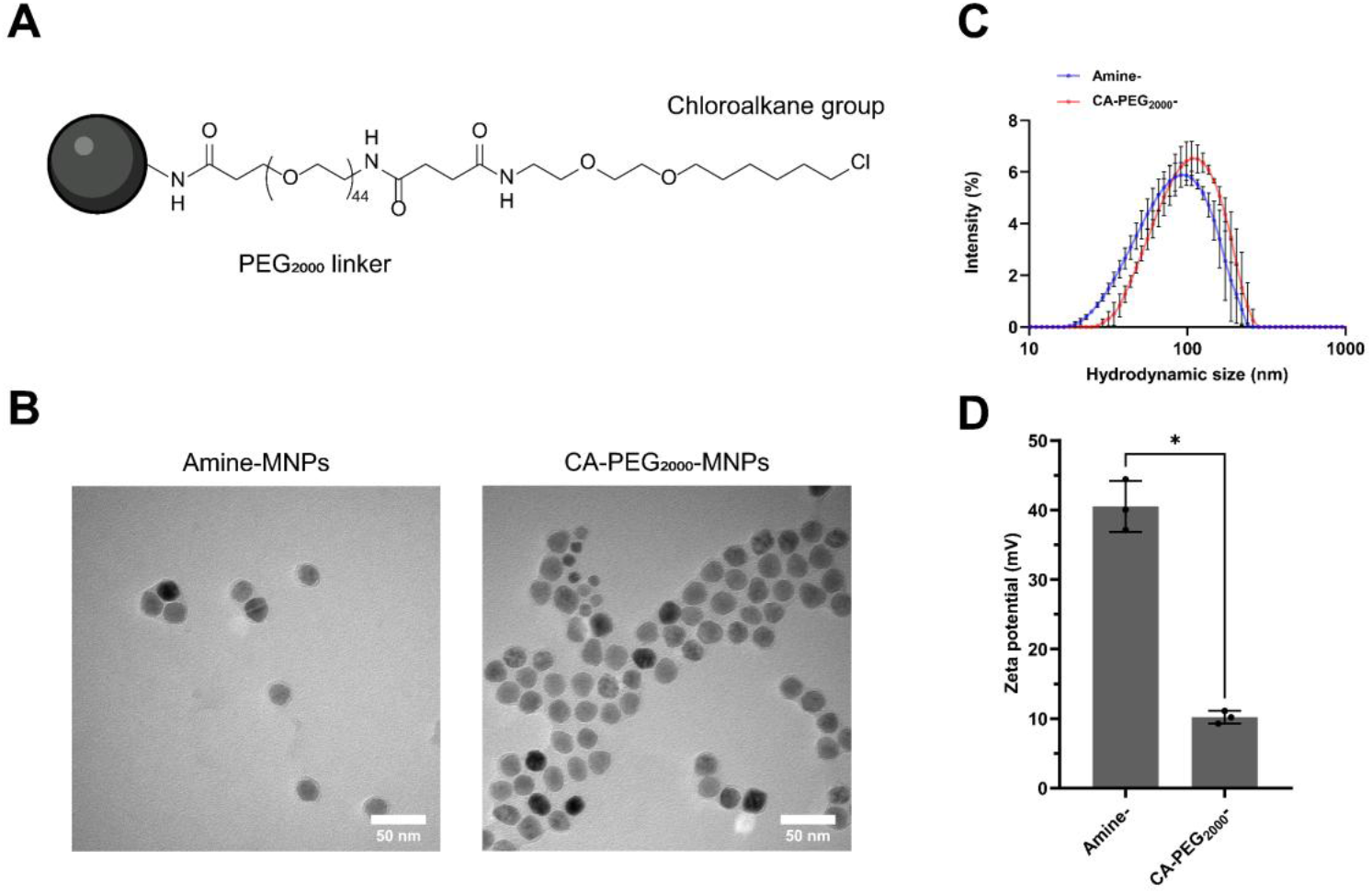
Chemical modification and characterization of magnetic nanoparticles (MNPs). **(A)** Schematic illustration of MNP modification with HaloTag ligands consisting of a PEG linker (2000 Da) terminated with a CA group. **(B)** TEM images of unmodified amine-functionalized MNPs and CA-PEG_2000_-MNPs. **(C)** Size distribution curves showing the hydrodynamic diameters of MNPs. **(D)** Zeta potential measurements of amine-MNPs and CA-PEG_2000_-MNPs. Data are presented as means ± SD, *n* = 3 biological replicates. Statistical significance was determined using the nonparametric Mann-Whitney *U* test. \**P* ≤ 0.05.

Starting from the initial amount of CA-PEG_2000_-NHS, we prepared serial dilutions at 1/2, 1/10, and 1/100 of the original concentration to modify the MNPs. When these particles were attached to *E. coli* (pWX118), we observed improved functionalization efficiency, as indicated by the formation of dark bacterial pellets indicating the attachment of MNPs (Fig. S11). Notably, at 1/100 of the original ligand amount, functionalization became robust, and magnetic torque-driven actuation under RMF could clearly be observed. We therefore prepared a series of MNPs functionalized with varying ligand concentrations for further characterization. Dynamic light scattering (DLS) analysis confirmed our hypothesis. When a large excess of CA-PEG_2000_ ligands were used, the hydrodynamic size of the MNPs remained almost the same as that of amine-terminated particles, suggesting that excess PEG molecules collapsed or cloaked together on the surface rather than extending outward (Fig. S12A and S12B). By contrast, when 1/100 or less of the ligand concentration was applied, the hydrodynamic size of all MNPs increased by 20-30 nm, indicating un-cloaked CA-PEG ligands. In addition, zeta potential measurements showed that all modified MNPs exhibited lower surface charge than amine-MNPs. When MNPs were functionalized with 1/100 or less of the ligand amount, the surface charge increased to 10–20 mV, as a higher fraction of positively charged amine groups remained (Fig. S12C).

Based on these results, we selected CA-MNPs functionalized with 1/100 of the initial ligand amount for detailed characterization. TEM imaging confirmed that both CA-MNPs and amine-MNPs exhibited similar morphology and were well dispersed without aggregation (**Fig. 4B**). DLS characterization of these particles showed that CA-MNPs had an average hydrodynamic diameter of 97.16 nm, larger than the 78.28 nm observed for amine-terminated MNPs (**Fig. 4C**), while both displayed a similar polydispersity of ∼20%. The average zeta potential of CA-MNPs was 10.19 mV, which is only about one quarter of the 40.53 mV measured for amine-MNPs (**Fig. 4D**). This indicates that the residual amine groups on CA-MNPs are unlikely to contribute significantly to electrostatic adsorption, thereby minimizing nonspecific binding to bacteria. Therefore, we finalized our protocol by using 1/100 of the original CA-PEG_2000_ ligand concentration, and the resulting CA-MNPs were used for subsequent experiments.

### Magnetic functionalization and characterization of HaloBots

After achieving a highly efficient and homogeneous HaloTag display on bacteria and optimizing the MNP modification process, we next used the CA-MNPs to functionalize HaloTag-displayed bacteria, generating HaloBots (**Fig. 5A**). Non-transformed *E. coli* was used as a control to assess whether electrostatic interactions between bacteria and MNPs could lead to nonspecific particle attachment. Magnetic functionalization was performed under a constant magnetic field to enhance the magnetic anisotropy of HaloBots, as described in our recent study^30^.

**Figure 5.**
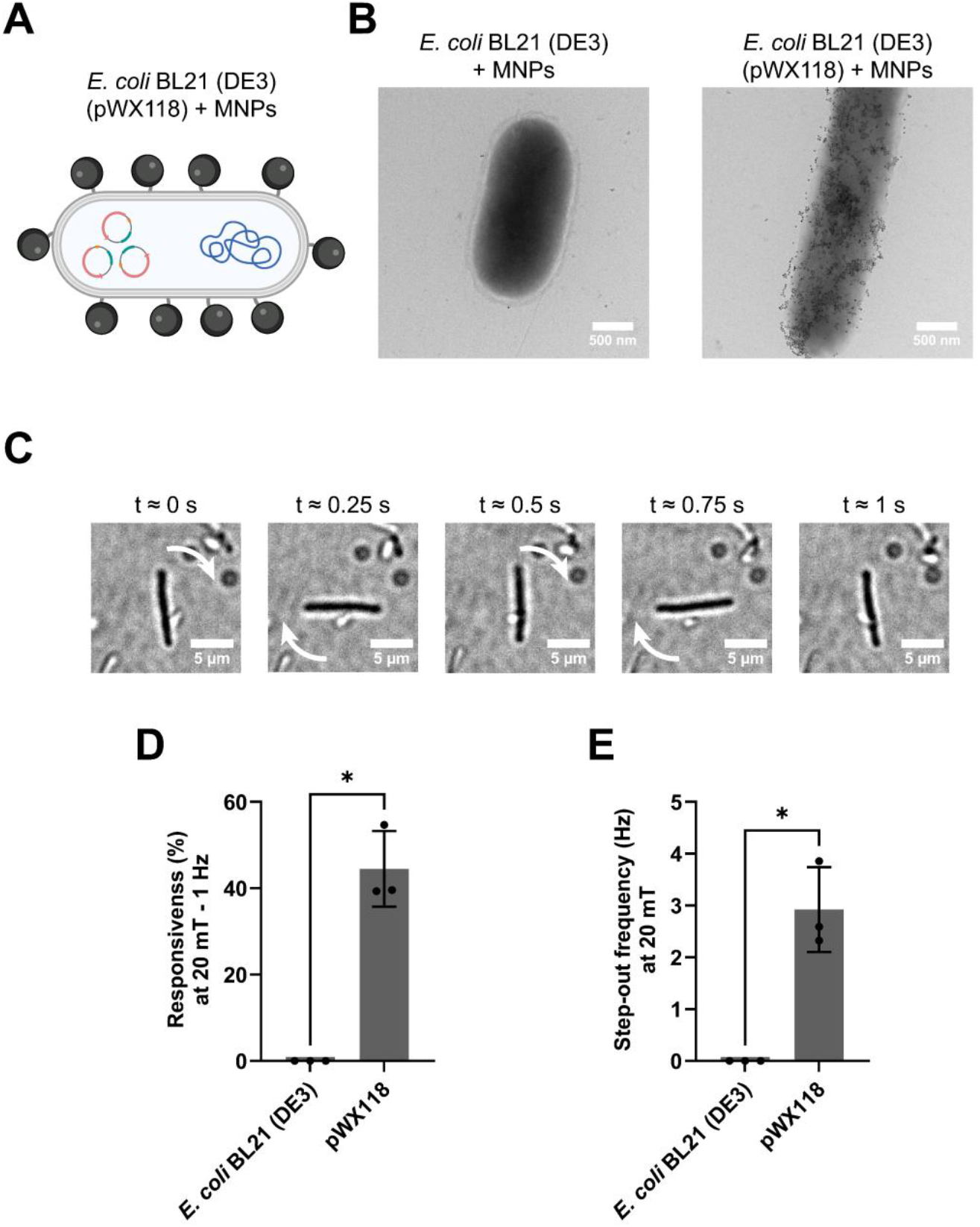
Magnetic functionalization and characterization of bacteria. **(A)** Schematic illustration of *E. coli* functionalized with MNPs. **(B)** TEM images of non-transformed *E. coli* and HaloTag-expressing *E. coli* after functionalization with MNPs. **(C)** Microscopy images showing the rotation of an *E. coli* cell actuated by a rotating magnetic field (RMF, 20 mT, 1 Hz). **(D)** Bacterial responsiveness (%) under a 20 mT, 1 Hz rotating magnetic field. **(E)** Step-out frequencies of bacteria under a 20 mT rotating magnetic field. Data are presented as means ± SD, *n* = 3 biological replicates. Statistical significance was determined using the nonparametric Mann-Whitney *U* test. \**P* ≤ 0.05.

Following functionalization, a clear distinction was observed: pWX118-transformed *E. coli* pellets remained dark, reflecting extensive CA-MNP conjugation, whereas control pellets, initially dark from weak electrostatic adsorption, turned nearly white after washing, indicating that most MNPs were removed (Fig. S13). TEM imaging further validated this difference (**Fig. 5B**). While non-transformed bacteria showed nearly no conjugation with MNPs, the engineered bacteria exhibited dense coverage of particles. A representative image sequence showcased that, under RMF, the functionalized *E. coli* rotated in alignment with the field (**Fig. 5C**). To investigate whether HaloTag display might alter the surface charge and thereby contribute to electrostatic adsorption between bacteria and MNPs, we measured the zeta potential of HaloTag-displayed bacteria and non-transformed bacteria. The results showed that the zeta potential of HaloTag-displayed bacteria was nearly identical to that of the non-transformed ones (Fig. S14), indicating that the robust magnetic functionalization observed in *E. coli* can be attributed to the specific covalent reaction between surface-displayed HaloTag and the CA ligands on the MNPs, rather than to nonspecific electrostatic interactions. We therefore concluded that HaloBots were successfully generated.

To characterize the magnetic responses of HaloBots, we actuated HaloBots (n=3) using a RMF at 20 mT, 1 Hz and analyzed the percentage (%) of responsive bacteria. In addition, the step-out frequency of HaloBots was analyzed to assess the maximum rotational speed at which HaloBots can remain synchronized with the driving field, providing insight into the magnitude of their magnetic moment. All experiments and analysis were performed using previously described methods.^30^ The results demonstrated that, on average, 44% of HaloBots met the responsiveness criterion (**Fig. 5D**), and the average step-out frequency was 2.9 Hz (**Fig. 5E**). In a clear contrast, non-transformed bacteria exhibited zero magnetic response.

These results demonstrate that HaloBots, generated by engineering *E. coli* to display HaloTag and covalently conjugated to ligand-modified MNPs, can be well controlled at swarm level using a magnetic field. Their magnetic response is robust and consistent, highlighting their potential as a promising magnetic bacterial microrobot.

## Discussion

This study highlights the potential of the self-labeling protein HaloTag for robust and specific magnetic functionalization of bacteria, addressing key challenges in bacterial microrobot development, including limited conjugated magnetic materials, inconsistent functionalization, weak binding, and nonspecific interactions. By engineering bacteria to display HaloTag on their outer surfaces and modifying MNPs with HaloTag ligands, we achieved efficient functionalization of HaloTag-displaying bacteria. The results demonstrate that successful functionalization depends on both genetic and chemical engineering. On the genetic side, the Lpp–OmpA scaffold and LF linker allow HaloTag to extend into the extracellular environment, facilitating conjugation with CA-MNPs. On the chemical side, both PEG linker length and ligand density are critical: long PEG linkers extend reactive CA groups but excessive PEGylation can reduce accessibility, whereas insufficient ligand density leaves exposed amines, increasing positive charge and causing nonspecific binding. By tuning ligand density, we achieved a balanced surface chemistry: MNPs remained slightly positively charged, allowing weak electrostatic attraction to enhance encounters with bacteria while maintaining HaloTag–CA covalent reaction as the dominant mechanism.

The successful functionalization of *E. coli* not only generated a new class of magnetic bacterial microrobots but also provides a versatile platform for broader applications. HaloTag-displayed bacteria can be functionalized with diverse cargoes, including photoacoustic particles for imaging, drug-loaded nanoliposomes for targeted delivery, or complex proteins that bacteria cannot natively express. Multiple cargo types can also be co-conjugated, enabling multimodal microrobots with combined functionalities. Additionally, the modularity of this approach opens several opportunities for future development. Different self-labeling proteins could be tested to expand functionalization strategies, and genetic or chemical components could be independently tuned to optimize cargo loading, binding strength, or responsiveness. Moreover, while we demonstrated the method in *E. coli*, alternative display scaffolds could be developed for other bacterial strains to broaden applicability.

Overall, HaloTag-based engineering provides a simple, modular, and robust strategy for bacterial microrobot functionalization. By integrating genetic surface display with covalent chemical conjugation, this strategy enables reliable magnetic control while remaining highly adaptable to diverse cargoes and functionalities. More broadly, this platform can be extended beyond magnetic systems to engineer bacterial microrobots with customizable functions.

## Materials and methods Materials

CA-COOH (Halo-PEG_2_-Suc) was purchased from Iris Biotech GmbH. NH_2_-PEG_2000_-COOH, O-(N-succinimidyl)-N,N,N′,N′-tetramethyluronium tetrafluoroborate (TSTU), DIPEA and solvents used for chemical synthesis were ordered from Merck. Amine-functionalized iron oxide nanoparticles were purchased from Ocean Nanotech (SHA25,5 mg/mL).

### Construction of plasmids

Plasmids used in this study were constructed using three different vectors. For protein expression in *E. coli*, we used pM965, containing the constitutive *rpsM* promoter, and pCDFDuet-1, containing the T7 promoter under control of the Lac operon, both kindly provided by Tim Keys (ETH Zurich). Additional plasmids were obtained from Addgene: pKB223, encoding the Lpp–OmpA–tether peptide (Addgene plasmid #170013, gift from Jeffrey Tabor), and pET-51b-HaloTag11, encoding HaloTag11 (Addgene plasmid #175519, gift from Kai Johnsson). All other linear DNA fragments required for cloning were synthesized by Twist Bioscience (USA), and oligonucleotides were synthesized by Microsynth AG (Switzerland). All plasmids were sequenced by Microsynth AG using Sanger sequencing.

To construct plasmid pWX100, the Lpp–OmpA fragment was amplified from pKB223 using oligonucleotides introducing EcoRI and SalI restriction sites. The HaloTag11 fragment was amplified from pET-51b-HaloTag11 with SalI and HindIII sites. The pM965 backbone was digested with EcoRI and HindIII, and the three DNA fragments were ligated using T4 DNA Ligase. All molecular cloning reagents were obtained from New England Biolabs (UK). To construct plasmid pWX104, the Lpp-OmpA-tether peptide fragment amplified from pKB223 introducing EcoRI and SalI restriction sites and inserted with the same HaloTag fragment between the EcoRI and HindIII restriction sites of pM965.

To construct plasmid pWX118, the HaloTag fragment was amplified using a forward primer containing two tandem repeats of the GGGGS peptide sequence and a reverse primer introducing SalI and HindIII restriction sites. The Lpp-OmpA-tether peptide fragment carrying EcoRI and SalI sites was inserted together with the HaloTag fragment into the EcoRI and HindIII restriction sites of the pCDFDuet-1 backbone.

### Bacteria engineering for surface display of HaloTag

To display HaloTag on the outer membrane of Gram-negative bacteria, *E. coli* BL21 (DE3) competent cells (C2527I, New England Biolabs) were transformed with 100 ng of plasmids pWX100, pWX104, or pWX118 by heat shock at 42 °C for 45 s. Cells were recovered in 950 µL LB medium and cultured overnight on LB agar plates containing 100 µg/mL ampicillin (for pWX100 and pWX104) or 50 µg/mL streptomycin (for pWX118).

To induce protein expression, fresh colonies of *E. coli* were inoculated into 5 mL of LB medium supplemented with the corresponding antibiotics and cultured overnight. The following day, 100 µL of the overnight culture was diluted into 10 mL fresh medium in a 50 mL baffled flask and shaken at 220 rpm until OD_600_ reached 0.5. Protein expression was induced by adding isopropyl β-D-thiogalactoside (IPTG) to a final concentration of 1 mM and incubated for 5 h at 37 °C. For constitutive expression or wild-type control, the bacteria were harvested when OD_600_ reached 0.6.

To purify the bacteria for better HaloTag display efficiency, bacterial colonies were streaked on fresh LB agar plates containing the antibiotics for two times. From the last plates, colonies were taken and cultured overnight in LB medium with the antibiotics. The next morning, bacteria were frozen with 15% of glycerol in a -80 °C freezer. For following experiments, these glycerol stocks were used as the starting culture.

### Synthesis of fluorescent HaloTag ligand

To synthesize the fluorescent HaloTag ligand CA-sCy5, 1 mg (1.35 μmol, 1 equiv.) of Sulfo-Cy5-amine was added to a solution of 0.6 mg of CA-NHS (1.35 μmol, 1 equiv.) and 1.2 μL (6.75 μmol, 5 equiv.) of DIPEA in 150 μL dry DMF. The solution was stirred for 1 h at room temperature. Post reaction the mixture was directly subjected to prep HPLC (Waters C18, 7 μm, 21.2 mm × 250 mm, solvent A: MQ with 0.1% TFA, solvent B: MeCN, at 20 mL/min, with a gradient of 20–60% B over 35 min) to give a blue lyophilizate (708 nmol, 52%).

### Labeling surface-displayed HaloTag using fluorescent ligands

Bacteria were chilled on ice for 15 min and the OD_600_ was measured again. The bacteria should be diluted if the OD_600_ is higher than 1.0 to make sure they are in the linear range. The volume of bacteria needed for labeling is calculated based on the following equation:

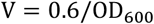

Specific volumes of bacteria were transferred into 2 mL Eppendorf tubes, washed three times with PBS by centrifugation at 5,000 × g for 2 min, and resuspended in 1 mL PBS. For labeling, CA-sCy5 solution was prepared by dissolving the solid dye in DMSO to a final concentration of 50 µM. 10 µL of this solution was added to 1 mL of bacterial suspension, resulting in a final dye concentration of 0.5 µM. The samples were incubated on an orbital shaker at 150 rpm for 1 h, followed by three additional PBS washes and resuspension for fluorescent imaging using an inverted spinning disk confocal microscope (Nikon Eclipse Ti2, equipped with a Yokogawa CSU-W1 unit and a Hamamatsu C13440-20CU Digital CMOS camera). A 100× objective was used for high resolution imaging with oil immersion.

### Synthesis of HaloTag ligand for MNP modification

The synthesis of HaloTag ligand CA-PEG_2000_-COOH includes two steps: 1) synthesis of CA-NHS by activation of CA-COOH with TSTU and DIPEA. 2) synthesis of CA-PEG_2000_-COOH by coupling CA-NHS to amine-PEG_2000_-COOH.

To synthesize CA-NHS, 64 mg (198 μmol, 1 equiv.) of Halo-PEG_2_-Suc and 89 mg (297 μmol, 1.5 equiv.) of TSTU were dissolved in 600 μL dry DMF under nitrogen atmosphere. 103 μL (594 μmol, 3 equiv.) of DIPEA were added and the solution was stirred for 1 h at room temperature. Post reaction 200 μL of AcOH, 1 mL acetonitrile and 1 mL of water were added and the solution was subjected to prep. HPLC ( Waters C18, 7 μm, 21.2 mm × 250 mm, solvent A: MQ with 0.1% TFA, solvent B: MeCN, at 20 mL/min, with a gradient of 20–70% B over 35 min) to give 52 mg (62%) of a colorless oil.

To synthesize CA-PEG_2000_-COOH, 40 mg (20 μmol, 1 equiv.) of NH2-PEG_2000_-COOH was added to a solution of 17 mg (40 μmol, 2 equiv.) of CA-NHS and 14 μL (80 μmol, 4 equiv.) of DIPEA in 1 mL dry DMF and the solution was stirred for 2 h at room temperature. Post reaction the reaction mixture was transferred to a dialysis tube (cutoff 1 kDa) and dialyzed 3 times against 1000 mL MQ water. The dialyzed solution was then lyophilized to give 30 mg (65%) of a white lyophilizate.

### Modification of MNPs with HaloTag ligands

Modification of MNPs with HaloTag ligands was carried out in two steps. The original protocol, which employed a large excess of HaloTag ligands, was as follows: First, 15 mg (6 μmol) of CA-PEG_2000_-COOH and 3 mg (10 μmol) of TSTU were dissolved in 200 µL of anhydrous DMSO in a 1.5 mL Eppendorf tube. Then, 3.5 µL (20 μmol) of DIPEA was added, and the mixture was incubated on an orbital shaker at 200 rpm for 1 h to yield CA-PEG_2000_-NHS. To modify MNPs, 200 µL of amine-functionalized iron oxide nanoparticles (5 mg/mL, estimated content of functional groups 10 nmol;^30^ Ocean Nanotech, SHA25) were dispersed in 800 µL PBS, after which the entire ligand solution was added. The suspension was incubated on an orbital shaker at 150 rpm for 3 h.

To prepare MNPs with reduced ligand density, the activated CA-PEG_2000_-NHS was diluted 100-fold in anhydrous DMSO, and 200 µL of the diluted solution was added to the MNP suspension for modification.

Excess ligands and byproducts were removed using M columns (Miltenyi Biotec) mounted on a µMACS Separator (Miltenyi Biotec) (Fig. S15). Columns were first rinsed with PBS using a plunger. The MNP suspension was diluted with 2 mL PBS to reduce the DMSO concentration and transferred to the column. After the liquid had passed through, the column was washed twice with 2 mL PBS and removed from the separator. MNPs were then eluted with 200 µL PBS, followed by an additional 100 µL to ensure complete elution.

To quantify the MNPs and determine the volume required for bacterial functionalization, their absorbance was measured using a microplate reader (Tecan Spark). The values were compared with a standard curve generated from amine-MNPs of known concentrations. Based on this, the MNP volume needed for conjugating 1 mL of bacteria was calculated, which is equivalent to approximately 5,000–10,000 particles per bacterium. The MNPs were stored at 4 °C.

### Characterization of MNPs

Dynamic light scattering (Litesizer, Anton Paar) was used to characterize MNPs. Briefly, 10 µL of CA-MNPs or amine-MNPs were diluted in 1 mL Milli-Q water, briefly sonicated, and vortexed. Hydrodynamic size was measured using disposable plastic cuvettes, while zeta potential was measured using an omega cuvette. After each measurement of zeta potential, the cuvette was thoroughly rinsed with Milli-Q water before reusing.

For TEM imaging, MNPs were prepared on carbon-coated grids. Grids were placed on a clean Kimtech wipe, and 10 µL of CA-MNPs or amine-MNPs, diluted in 500 µL Milli-Q water, were applied dropwise until the suspension was completely used. The grids were air-dried for 1 h and subsequently vacuum-dried overnight.

### Magnetic functionalization and actuation of bacteria

Bacteria were washed and resuspended in 1 mL of PBS in 2 mL Eppendorf tubes, as described above. CA-MNPs were retrieved from storage and briefly sonicated. Based on the calculated volume required for functionalization, MNPs were added to the bacterial suspensions in 10 µL increments, with immediate vortexing after each addition to minimize aggregation. Samples were incubated on an orbital shaker at 150 rpm for 1 h under a constant magnetic field (30 mT) genereted a Lee-Whiting coil,^30^ followed by three additional PBS washes.

For magnetic actuation, the bacteria were resuspended in Milli-Q water to reduce electrostatic adsorption between the cells and glass slides. 10 µL of the bacterial suspension were placed on a glass slide, and actuation was performed using a small-scale arbitrary magnetic field generator (MFG-100-I, MagnebotiX, Switzerland) mounted on a spinning-disk confocal microscope for real-time video recording. For a quick observation of the magnetic response, bacteria were actuated by an in-plane rotating magnetic field (RMF, 20 mT, 1 Hz) for 10 s. Videos were acquired and processed with Fiji software to export real-time AVI files.

### Characterization of magnetic responsiveness and step-out frequency

To quantify magnetic responsiveness and determine the step-out frequency, bacteria were imaged using a 60× objective during actuation with a sweeping in-plane rotating magnetic field (RMF; 20–0 mT, 1 Hz) for 20 s. Recorded videos were analyzed in Fiji and subjected to analysis employing the Feature-Assisted Segmenter/Tracker algorithm (FAST), which enabled segmentation, feature extractions, and tracking of 30–100 bacteria per video. Data from multiple videos were combined to ensure that at least 150 bacteria were analyzed per sample. The fraction of responsive bacteria was then calculated using an in-house MATLAB script, classifying a bacterium as responsive when its rotational frequency exceeded 90% of the applied RMF frequency. The step-out frequency of the population was defined as the RMF frequency at which bacterial responsiveness decreased to 50%, as described in our previous study.^30^

### Transmission electron microscopy (TEM) imaging

To prepare bacterial samples for TEM imaging, carbon-coated grids were first treated with air plasma. Briefly, fresh carbon grids were placed in a clean glass vial with the carbon-coated side facing upward. The vial was placed in a Diener Zepto Plasma Cleaner and treated with plasma for 12 s. Bacteria functionalized with MNPs were centrifuged and resuspended in 500 µL Milli-Q water to concentrate the sample. 10 µL of the suspension were applied to each grid and allowed to adsorb for 30 s. Excess liquid was removed using Kimtech wipes, and the deposition step was repeated twice. The grids were then air-dried for 1 h and subsequently vacuum-dried overnight.

### Statistical analysis

All data are presented as mean ± standard deviation (SD). Graphs were generated using GraphPad Prism 10 software. For comparisons between two groups, the nonparametric Mann-Whitney U test was used. Significance levels were indicated as not significant (ns) P > 0.05, *P ≤ 0.05. Experiments were performed in biological triplicates (*n* = 3).

### Declaration of AI-assisted technologies in the writing process

During the preparation of this manuscript, the AI language model ChatGPT (version GPT 5 mini) was used to improve the clarity and readability of certain portions of the text. All authors subsequently reviewed and edited the content as necessary and take full responsibility for the accuracy and integrity of the work.

## Acknowledgments

This study was funded by the Swiss National Science Foundation (SNSF) (10001430). The authors gratefully acknowledge Ines Oberhuber for her support with TEM imaging. We also thank Tim Keys for his suggestions on the molecular cloning of plasmids. Additionally, early experimental work on this project was supported by Anagh Sinha as part of his master’s thesis research.

## Author contributions

X.W. and P.P. contributed equally to the work. S.S., P.P. and X.W. conceived the idea and designed the experiments. X.W. carried out the genetic engineering and magnetic functionalization of bacteria. P.P. performed the chemical synthesis and modification of magnetic nanoparticles. X.W., P.P., E.T., and S.S. contributed to interpreting the results. X.W. took the lead in writing the manuscript. All authors contributed to the final writing of the manuscript.

## References: Uncategorized References

1 P. A. Yang, Y. X. Han, C. Y. Mo, J. F. Luo, M. J. Shou, X. L. Gong, and Z. H. Zhou, Chemical Engineering Journal 520 (2025).

2 S. L. Zhu, Y. F. Cheng, J. Wang, G. L. Liu, T. T. Luo, X. J. Li, S. L. Yang, and R. H. Yang, Acta Biomaterialia 169, 88 (2023).

3 M. Sitti and D. S. Wiersma, Advanced Materials 32 (20) (2020).

4 E. Totter, E. von Einsiedel, L. Regazzoni, and S. Schuerle, Advanced Drug Delivery Reviews 221 (2025).

5 T. Gwisai, N. Mirkhani, M. G. Christiansen, T. T. Nguyen, V. Ling, and S. Schuerle, Sci Robot 7 (71) (2022).

6 X. H. Yan, Q. Zhou, M. Vincent, Y. Deng, J. F. Yu, J. B. Xu, T. T. Xu, T. Tang, L. M. Bian, Y. X. J. Wang, K. Kostarelos, and L. Zhang, Sci Robot 2 (12) (2017).

7 M. B. Akolpoglu, Y. Alapan, N. O. Dogan, S. F. Baltaci, O. Yasa, G. A. Tural, and M. Sitti, Science Advances 8 (28) (2022).

8 C. K. Schmidt, M. Medina-Sánchez, R. J. Edmondson, and O. G. Schmidt, Nature Communications 11 (1) (2020).

9 H. T. Chen, Y. Z. Li, Y. J. Wang, P. Ning, Y. J. Shen, X. Y. Wei, Q. S. Feng, Y. L. Liu, Z. G. Li, C. Xu, S. Y. Huang, C. J. Deng, P. Wang, and Y. Cheng, Acs Nano 16 (4), 6118 (2022).

10 B. Wang, Y. W. Qin, J. Liu, Z. F. Zhang, W. H. Li, G. J. Pu, Z. Yuanhe, X. Gui, and M. Q. Chu, Acs Applied Materials & Interfaces 15 (2), 2747 (2023).

11 L. Zhang, Z. Chen, H. Ran, M. Azechi, X. Yang, W. Huang, H. Tian, L. Shen, F. Peng, and Y. Tu, Nat Commun 16 (1), 7856 (2025).

12 H. Chen, T. Zhou, S. Li, J. Feng, W. Li, L. Li, X. Zhou, M. Wang, F. Li, X. Zhao, and L. Ren, ACS Appl Mater Interfaces 15 (41), 47930 (2023).

13 A. Roda, L. Cevenini, S. Borg, E. Michelini, M. M. Calabretta, and D. Schüler, Lab on a Chip 13 (24), 4881 (2013).

14 D. Faivre and D. Schüler, Chemical Reviews 108 (11), 4875 (2008).

15 M. Aubry, W. A. Wang, Y. Guyodo, E. Delacou, J. M. Guigner, O. Espeli, A. Lebreton, F. Guyot, and Z. Gueroui, Acs Synthetic Biology 9 (11), 3030 (2020).

16 X. L. Liu, P. A. Lopez, T. W. Giessen, M. Giles, J. C. Way, and P. A. Silver, Scientific Reports 6 (2016).

17 M. V. Dziuba, A. Paulus, L. Schramm, R. P. Awal, M. Pósfai, C. L. Monteil, S. Fouteau, R. Uebe, and D. Schüler, Isme Journal 17 (3), 326 (2023).

18 M. V. Dziuba, F. D. Müller, M. Pósfai, and D. Schüler, Nature Nanotechnology 19 (1) (2024).

19 D. Y. Zhang, R. F. Fakhrullin, M. Özmen, H. Wang, J. Wang, V. N. Paunov, G. H. Li, and W. E. Huang, Microbial Biotechnology 4 (1), 89 (2011).

20 M. Martín, F. Carmona, R. Cuesta, D. Rondón, N. Gálvez, and J. M. Domínguez-Vera, Advanced Functional Materials 24 (23), 3489 (2014).

21 X. T. Ma, X. L. Liang, Y. Li, Q. Q. Feng, K. M. Cheng, N. N. Ma, F. Zhu, X. J. Guo, Y. L. Yue, G. N. Liu, T. J. Zhang, J. Liang, L. Ren, X. Zhao, and G. J. Nie, Nature Communications 14 (1) (2023).

22 M. Gorohovs and Y. Dekhtyar, Molecules 30 (15) (2025).

23 M. Mahmoudi, A. M. Abdelmonem, S. Behzadi, J. H. Clement, S. Dutz, M. R. Ejtehadi, R. Hartmann, K. Kantner, U. Linne, P. Maffre, S. Metzler, M. K. Moghadam, C. Pfeiffer, M. Rezaei, P. Ruiz-Lozano, V. Serpooshan, M. A. Shokrgozar, G. U. Nienhaus, and W. J. Parak, Acs Nano 7 (8), 6555 (2013).

24 S. Pasche, J. Vörös, H. J. Griesser, N. D. Spencer, and M. Textor, Journal of Physical Chemistry B 109 (37), 17545 (2005).

25 A. H. A. Balzer and C. B. Whitehurst, Current Issues in Molecular Biology 45 (11), 8733 (2023).

26 K. Boeneman, J. R. Deschamps, S. Buckhout-White, D. E. Prasuhn, J. B. Blanco-Canosa, P. E. Dawson, M. H. Stewart, K. Susumu, E. R. Goldman, M. Ancona, and I. L. Medintz, Acs Nano 4 (12), 7253 (2010).

27 K. W. Yong, D. Yuen, M. Z. Chen, C. J. H. Porter, and A. P. R. Johnston, Nano Letters 19 (3), 1827 (2019).

28 R. E. Bird, S. A. Lemmel, X. Yu, and Q. A. Zhou, Bioconjugate Chemistry 32 (12), 2457 (2021).

29 Q. Y. Zheng and P. Chang, Israel Journal of Chemistry 63 (1-2) (2023).

30 L. Regazzoni, E. Totter, S. Menghini, I. Oberhuber, F. Li, C. Forbrigger, P. Poc, X. Wang, C. Schirmer, P. Wagner, M. G. Christiansen, and S. Schuerle, Cell Reports Physical Science in press (2025).

31 G. V. Los, L. P. Encell, M. G. McDougall, D. D. Hartzell, N. Karassina, C. Zimprich, M. G. Wood, R. Learish, R. F. Ohana, M. Urh, D. Simpson, J. Mendez, K. Zimmerman, P. Otto, G. Vidugiris, J. Zhu, A. Darzins, D. H. Klaubert, R. F. Bulleit, and K. V. Wood, ACS Chem Biol 3 (6), 373 (2008).

32 C. G. England, H. M. Luo, and W. B. Cai, Bioconjugate Chemistry 26 (6), 975 (2015).

33 S. Engin, D. Fichtner, D. Wedlich, and L. Fruk, Current Pharmaceutical Design 19 (30), 5443 (2013).

34 K. Kolberg, C. Puettmann, A. Pardo, J. Fitting, and S. Barth, Current Pharmaceutical Design 19 (30), 5406 (2013).

35 S. Hoehnel and M. P. Lutolf, Bioconjugate Chemistry 26 (8), 1678 (2015).

36 S. Kühn, V. Nasufovic, J. Wilhelm, J. Kompa, E. M. F. de Lange, Y. H. Lin, C. Egoldt, J. Fischer, A. Lennoi, M. Tarnawski, J. Reinstein, R. Vlijm, J. Hiblot, and K. Johnsson, Nat Chem Biol 21 (11), 1754 (2025).

37 R. Minner-Meinen, J. N. Weber, A. Albrecht, R. Matis, M. Behnecke, C. Tietge, S. Frank, J. Schulze, H. Buschmann, P. J. Walla, R. R. Mendel, R. Hänsch, and D. Kaufholdt, Plant Commun 2 (5) (2021).

38 D. L. Buckley, K. Raina, N. Darricarrere, J. Hines, J. L. Gustafson, I. E. Smith, A. H. Miah, J. D. Harling, and C. M. Crews, ACS Chem Biol 10 (8), 1831 (2015).

39 J. Wilhelm, S. Kühn, M. Tarnawski, G. Gotthard, J. Tünnermann, T. Tänzer, J. Karpenko, N. Mertes, L. Xue, U. Uhrig, J. Reinstein, J. Hiblot, and K. Johnsson, Biochemistry 60 (33), 2560 (2021).

40 S. M. Marques, M. Slanska, K. Chmelova, R. Chaloupkova, M. Marek, S. Clark, J. Damborsky, E. T. Kool, D. Bodnar, and Z. Prokop, Jacs Au 2 (6), 1324 (2022).

41 A. Cook, F. Walterspiel, and C. Deo, Chembiochem 24 (12) (2023).

42 A. Pulsipher, M. E. Griffin, S. E. Stone, and L. C. Hsieh-Wilson, Angewandte Chemie-International Edition 54 (5), 1466 (2015).

43 T. Berki, A. Bakunts, D. Duret, L. Fabre, C. Ladavière, A. Orsi, M. T. Charreyre, A. Raimondi, E. van Anken, and A. Favier, Acs Omega 4 (7), 12841 (2019).

44 P. Xu, S. M. Zhong, Y. F. Wei, X. X. Duan, M. Zhang, W. Shen, Y. Ma, and Y. H. Zhang, Acs Nano 18 (32), 21433 (2024).

45 C. F. Earhart, Applications of Chimeric Genes and Hybrid Proteins, Pt A 326, 506 (2000).

46 P. H. Bessette, F. Åslund, J. Beckwith, and G. Georgiou, Proceedings of the National Academy of Sciences of the United States of America 96 (24), 13703 (1999).

47 C. Stathopoulos, G. Georgiou, and C. F. Earhart, Applied Microbiology and Biotechnology 45 (1-2), 112 (1996).

48 K. R. Brink, M. G. Hunt, A. M. Mu, K. Groszman, K. V. Hoang, K. P. Lorch, B. H. Pogostin, J. S. Gunn, and J. J. Tabor, Nat Chem Biol 19 (4), 451 (2023).

49 F. Khan, G. J. Jeong, N. Tabassum, A. Mishra, and Y. M. Kim, Applied Microbiology and Biotechnology 106 (18), 5835 (2022).

50 Q. G. Xu, L. M. Ensign, N. J. Boylan, A. Schön, X. Q. Gong, J. C. Yang, N. W. Lamb, S. T. Cai, T. Yu, E. Freire, and J. Hanes, Acs Nano 9 (9), 9217 (2015).

