## Supplementary Materials for "A modular HaloTag platform for engineering magnetically responsive bacterial microrobots"

**Supplementary Tables and Figures**

**Table S1. Plasmids used in this study and construction methods of plasmids.**

| Plasmids | Description | Reference |
|---|---|---|
| pM965 | Plasmid encoding constitutive expression unit of GFP (P*rpsM*-GFPmut2), and Ampicillin resistance. | (*180*) |
| pCDFDuet-1 | Plasmid encoding IPTG-inducible expression unit (T7-lacO) with dual multiple cloning sites (MCS1, MCS2) and streptomycin/spectinomycin resistance for *E. coli*. | Novagen, Merck |
| pKB223 | Plasmid encoding IPTG-inducible surface display of C18G-mNeonGreen via Lpp–OmpA tether (P*tac* promoter) for *E. coli*, and Ampicillin resistance. | Addgene plasmid # 170013 |
| pET-51b-HaloTag11 | Plasmid encoding IPTG-inducible expression of HaloTag11 (N-terminal His-tag) in *E. coli*, and Ampicillin resistance. | Addgene plasmid # 175519 |
| pWX100 | Plasmid encoding constitutive expression unit (P*rpsM*-Lpp-OmpA-HaloTag11) for *E. coli*, and Ampicillin resistance.<br><br>The Lpp-OmpA fragment was PCR amplified from pKB223 using oligonucleotides 5’-CCGGAATTCATGAAAGCTACTAAACTGGTACTGG-3’ and 5’- GCGTCGACGTTGTCCGGACGAGT-3’. The HaloTag 11 fragment was PCR amplified from pET-51b-HaloTag11 using oligonucleotides 5’-GCGTCGACGGTGGAGGCGGTTCAATCGGTACTGGCTTTCCATTCGAC-3’ and 5’-CCCAAGCTTTTAAATCTCCAGAGTAGACAGCCA-3’. The pM965 backbone was linearized by digesting with EcoRI and HindIII, and the two DNA fragments were ligated together into the backbone to generate plasmid pWX100. | This work |

| | | |
|---|---|---|
| pWX104 | Plasmid encoding constitutive expression unit (P*rpsM*-Lpp-OmpA-tether peptide-HaloTag11) for *E. coli*, and Ampicillin resistance.<br><br>The Lpp-OmpA-tether peptide fragment was PCR amplified from pKB223 using oligonucleotides 5'-CCGGAATTCATGAAAGCTACTAAACTGGTACTGG-3' and 5'-GCGTCGACGGTTCCTCCGATACCCG-3'. The HaloTag 11 fragment was PCR amplified from pET-51b-HaloTag11 using oligonucleotides 5'-GCGTCGACGGTGGAGGCGGTTCAATCGGTACTGGCTTTCCATTCGAC-3' and 5'-CCCAAGCTTTTAAATCTCCAGAGTAGACAGCCA-3'. The pM965 backbone was linearized by digesting with EcoRI and HindIII, and the two DNA fragments were ligated together into the backbone to generate plasmid pWX104. | This work |
| pWX118 | Plasmid encoding an IPTG-inducible expression unit (T7-Lpp-OmpA-tether peptide-$(GGGGS)_2$-HaloTag11) for *E. coli*, and streptomycin/spectinomycin resistance.<br><br>The Lpp-OmpA-tether peptide fragment was PCR amplified from pKB223 using oligonucleotides 5'-CCGGAATTCATGAAAGCTACTAAACTGGTACTGG-3' and 5'-GCGTCGACGGTTCCTCCGATACCCG-3'. The $(GGGGS)_2$-HaloTag 11 fragment was PCR amplified from pET-51b-HaloTag11 using oligonucleotides 5'-GCGTCGACGGTGGAGGCGGTTCAGGCGGAGGTGGCTCTATCGGTACTGGCTTTCCATTCGAC-3' and 5'-CCCAAGCTTTTAAATCTCCAGAGTAGACAGCCA-3'. The pCDFDuet-1 backbone was linearized by | This work |

| | |
|---|---|
| | digesting with EcoRI and HindIII, and the two DNA fragments were ligated together into the backbone to generate plasmid pWX118. |

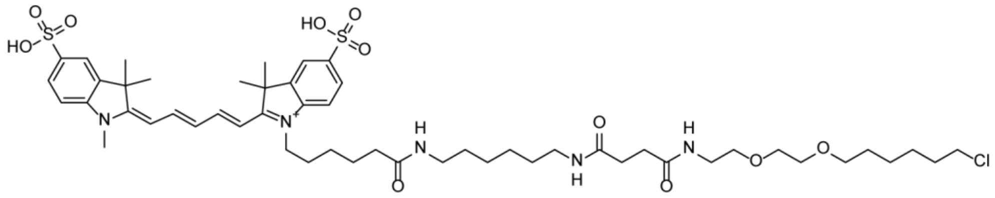

**Figure S1. Chemical structure of CA-sCy5.**

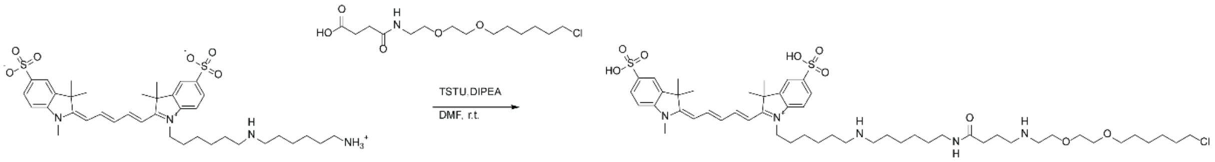

**Figure S2. Synthesis scheme of CA-sCy5.** The synthesis of CA-sCy5 was carried out in a one-step reaction by coupling Sulfo-Cy5-amine with CA-NHS using TSTU and DIPEA, thereby attaching the CA group to Sulfo-Cy5.

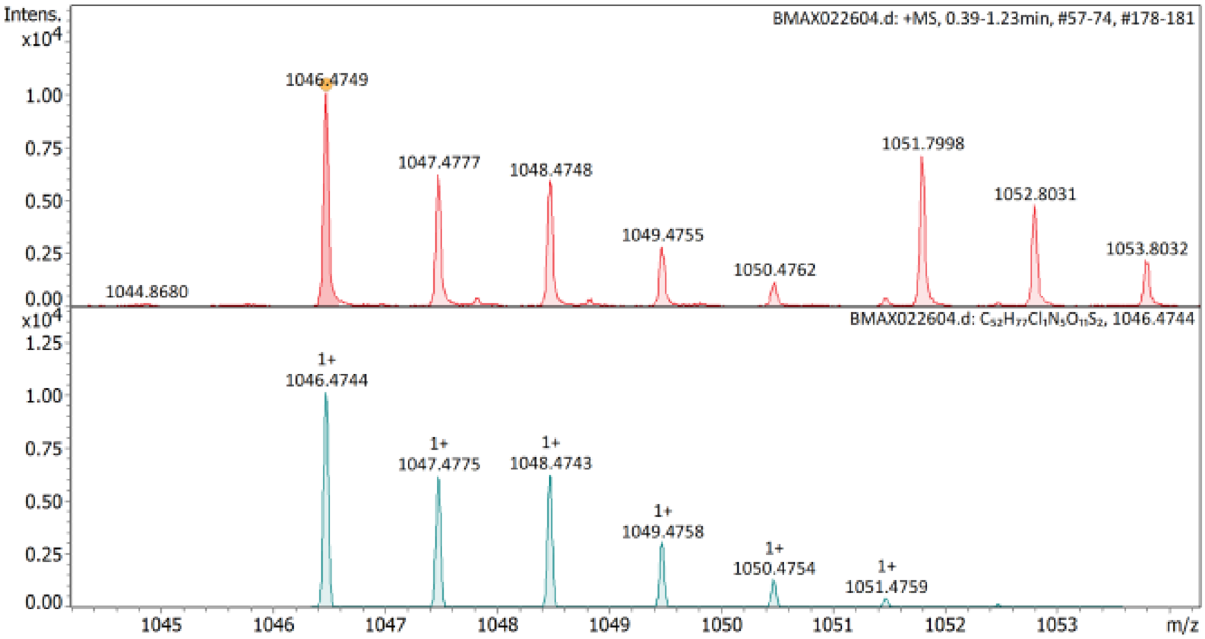

**Figure S3. High-resolution mass spectrometry (HR-MS) of CA-sCy5:** HRMS (ESI) m/z: calc. for $[C52H77ClN5O11S2]^{+}$: 1046.4744, found: 1046.4749

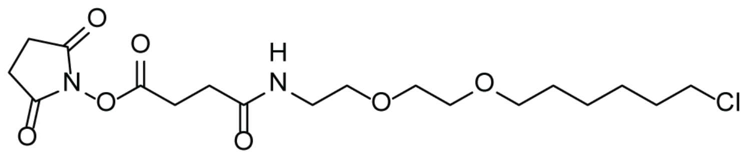

**Figure S4. Chemical structure of CA-NHS.**

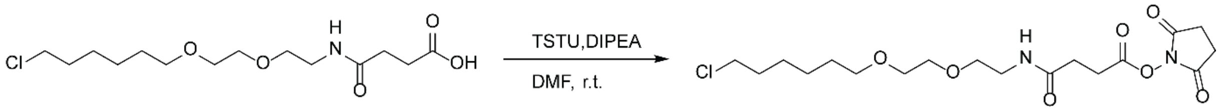

**Figure S5. Synthesis scheme of CA-NHS.** The synthesis of CA-NHS was carried out by coupling CA-COOH with NHS groups using TSTU and DIPEA.

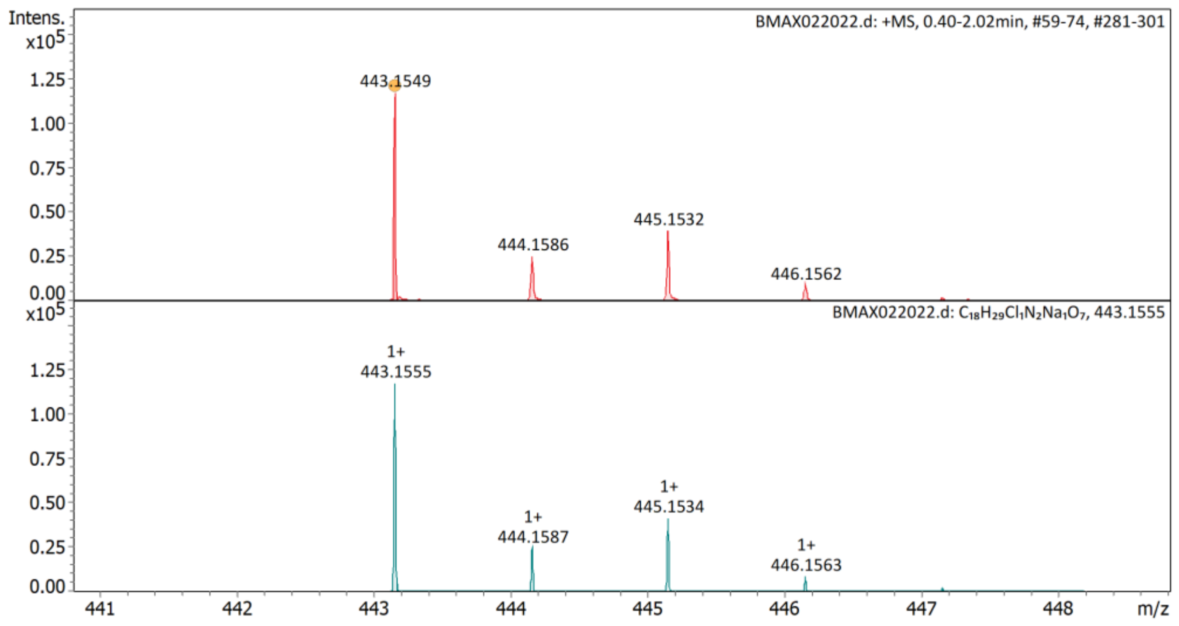

**Figure S6. High-resolution mass spectrometry (HR-MS) of CA-NHS:** HR-MS (ESI) m/z: calc. for [C18H29ClN2NaO7]$^{+}$: 443.1561, found: 443.1549

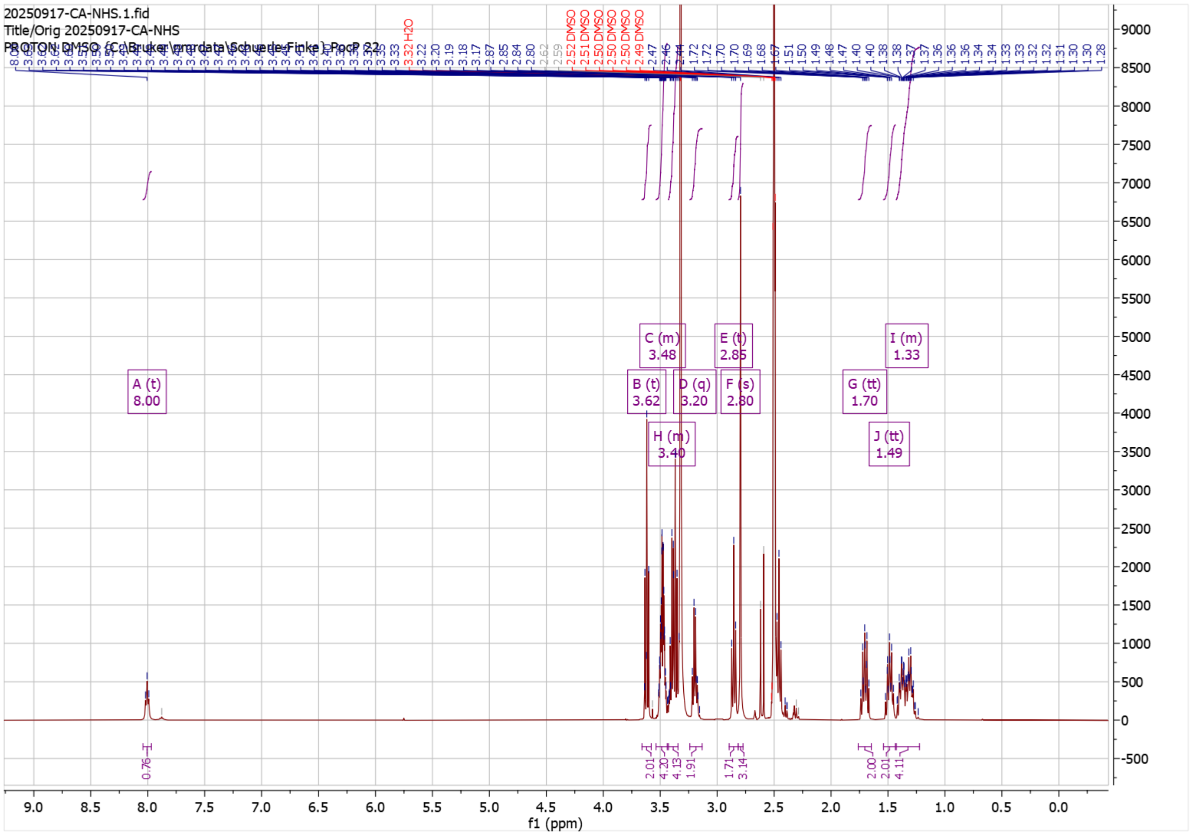

**Figure S7. ¹H-NMR spectrum of CA-NHS:** ¹H NMR (400 MHz, DMSO) δ 8.00 (t, *J* = 5.6 Hz, 1H), 3.62 (t, *J* = 6.6 Hz, 2H), 3.54 – 3.44 (m, 4H), 3.43 – 3.34 (m, 4H), 3.20 (q, *J* = 5.9 Hz, 2H), 2.85 (t, *J* = 6.9 Hz, 2H), 2.80 (s, 3H), 1.70 (tt, *J* = 7.8, 6.6 Hz, 2H), 1.49 (tt, *J* = 9.4, 6.4 Hz, 2H), 1.43 – 1.22 (m, 4H).

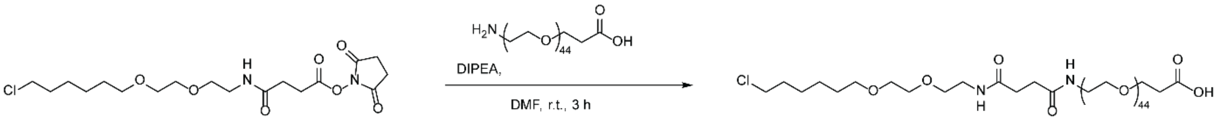

**Figure S8. Synthesis scheme of CA-PEG$_{2000}$-COOH.** The synthesis of CA-PEG$_{2000}$-COOH was carried out by coupling CA-NHS to amine-PEG$_{2000}$-COOH.

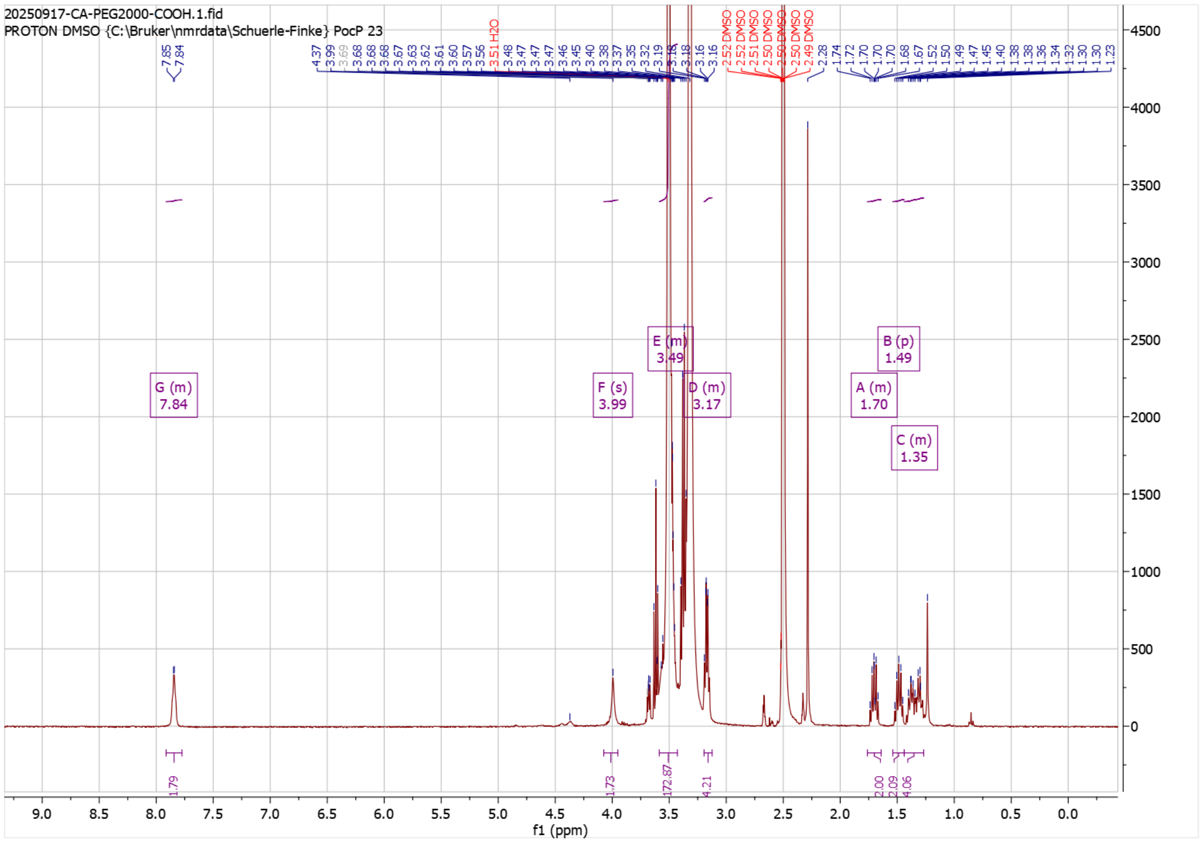

**Figure S9. $^1$H-NMR spectrum of CA-PEG$_{2000}$-COOH:** $^1$H NMR (400 MHz, DMSO) δ 7.91 – 7.77 (m, 2H), 3.99 (s, 2H), 3.59 – 3.43 (m, 173H), 3.19 – 3.12 (m, 4H), 1.76 – 1.64 (m, 2H), 1.49 (p, 2H), 1.44 – 1.27 (m, 4H).

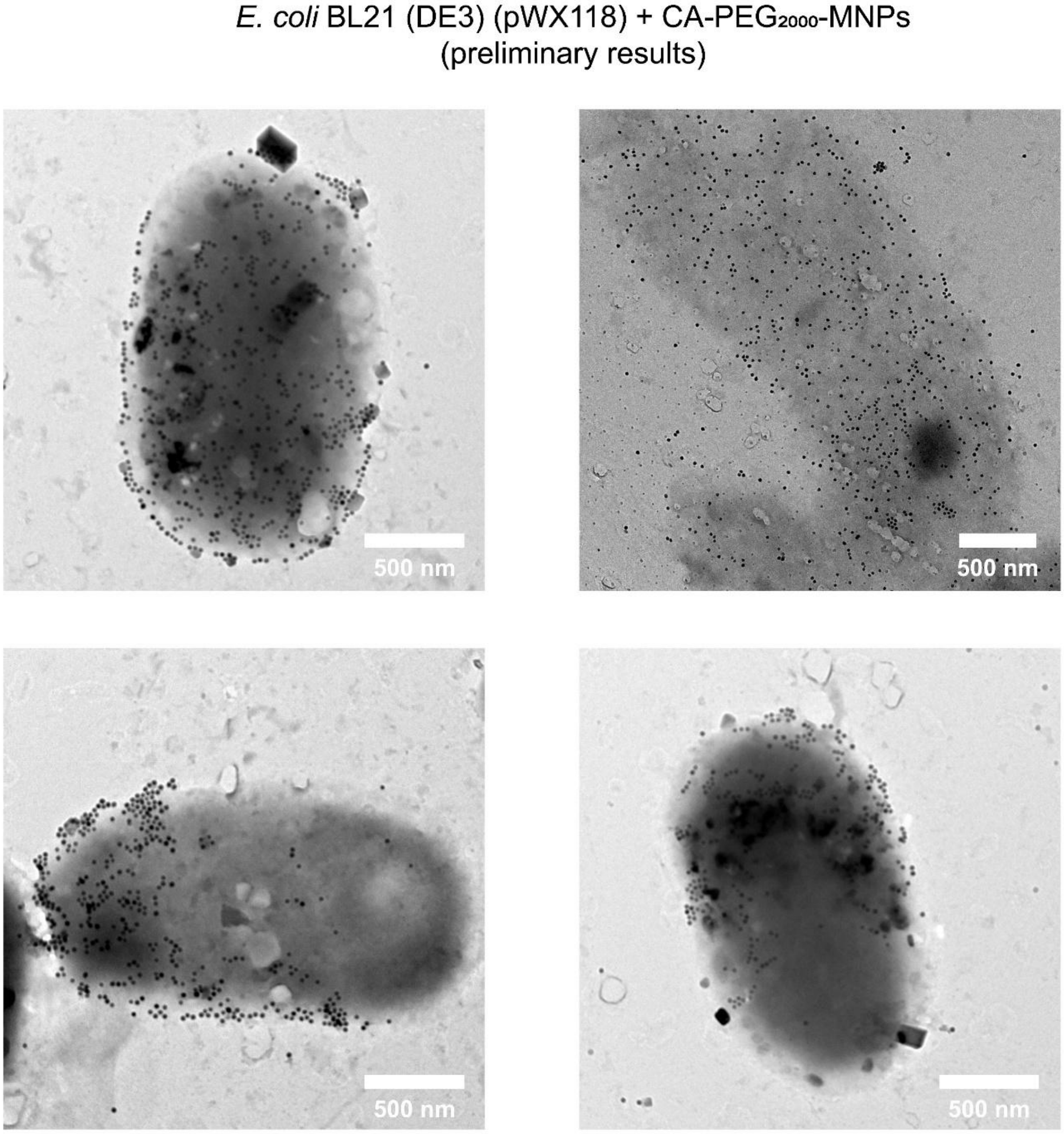

**Figure S10. Preliminary results of TEM images of *E. coli* BL21 (DE3) transformed with pWX118 and functionalized with CA-PEG$_{2000}$-MNPs (original ligand density).**

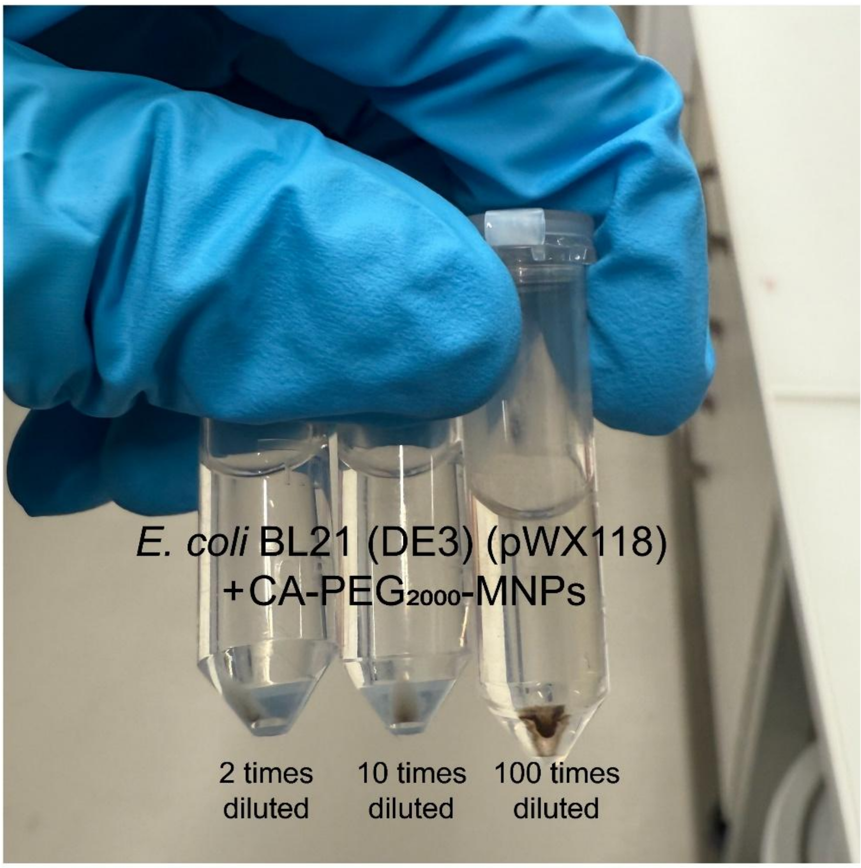

**Figure S11. Images of bacterial pellets of *E. coli* BL21 (DE3) transformed with pWX118 and functionalized with CA-PEG$_{2000}$-MNPs (ligand diluted 2, 10, and 100 times).**

A

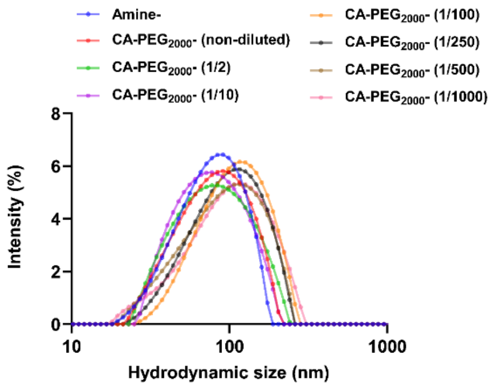

C

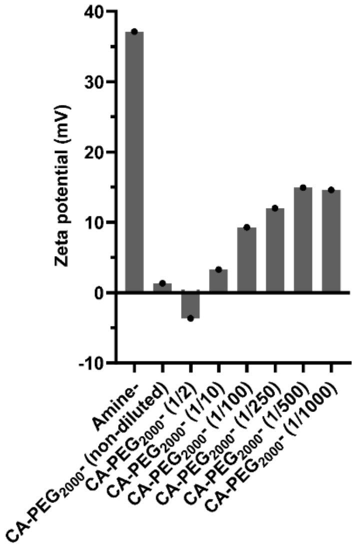

B

| Modification | Amine- | CA-PEG$_{2000}$- (non-diluted) | CA-PEG$_{2000}$- (1/2) | CA-PEG$_{2000}$- (1/10) | CA-PEG$_{2000}$- (1/100) | CA-PEG$_{2000}$- (1/250) | CA-PEG$_{2000}$- (1/500) | CA-PEG$_{2000}$- (1/1000) |
|---|---|---|---|---|---|---|---|---|
| Mean size (nm) | 74.98 | 76.05 | 75.12 | 71.90 | 100.22 | 91.55 | 88.05 | 89.36 |
| Polydispersity (%) | 24.54 | 22.29 | 23.54 | 21.04 | 22.82 | 23.69 | 24.78 | 24.76 |

**Figure S12. Hydrodynamic size and zeta potential of amine-MNPs and CA-PEG$_{2000}$-MNPs (serial ligand dilutions).**

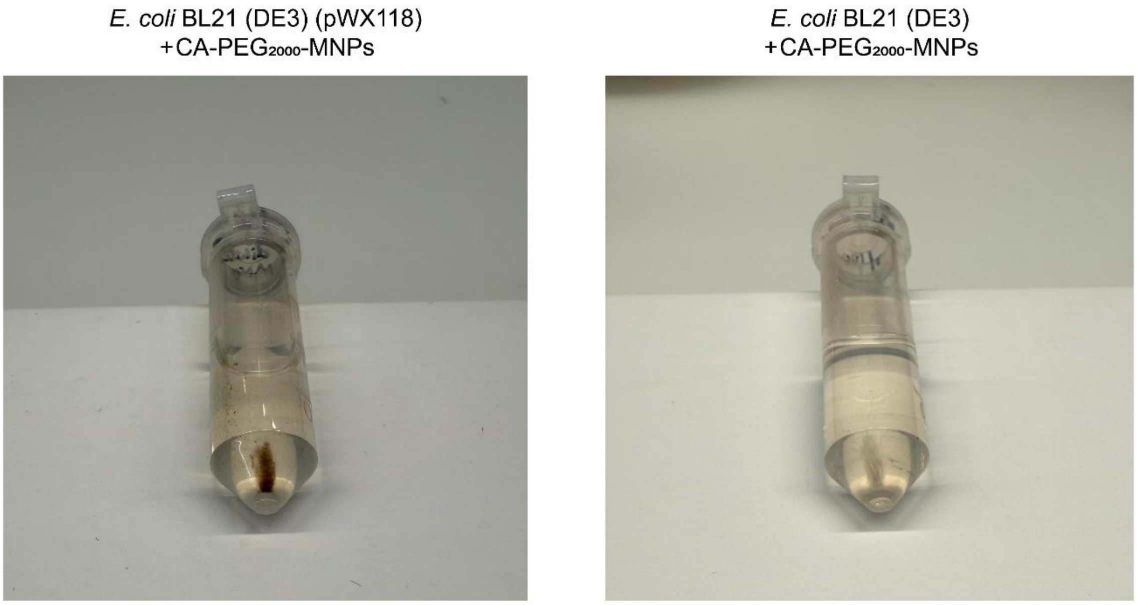

**Figure S13. Images of bacterial pellets of *E. coli* BL21 (DE3) transformed with pWX118 and non-transformed *E. coli* functionalized with CA-PEG$_{2000}$-MNPs (ligand diluted 100 times).**

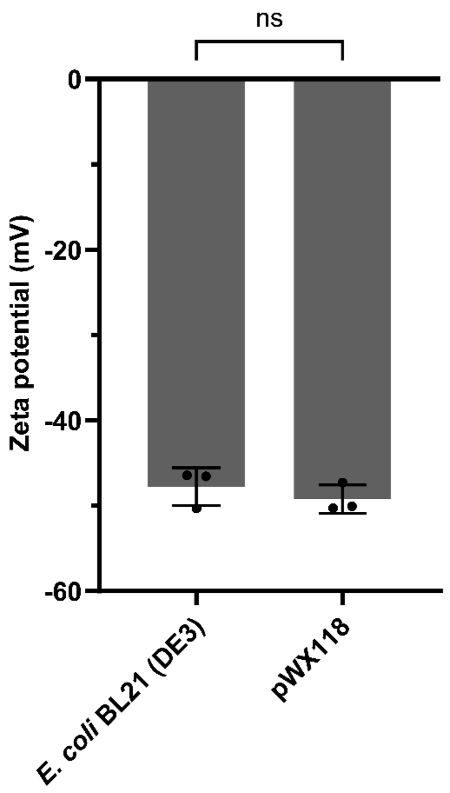

**Figure S14. Zeta potential of non-transformed *E. coli* BL21 (DE3) and *E. coli* transformed with pWX118.** Data are presented as means ± SD, $n = 3$ biological replicates. Statistical significance was determined using the nonparametric Mann-Whitney *U* test. Significance levels were indicated as not significant (ns) P > 0.05.

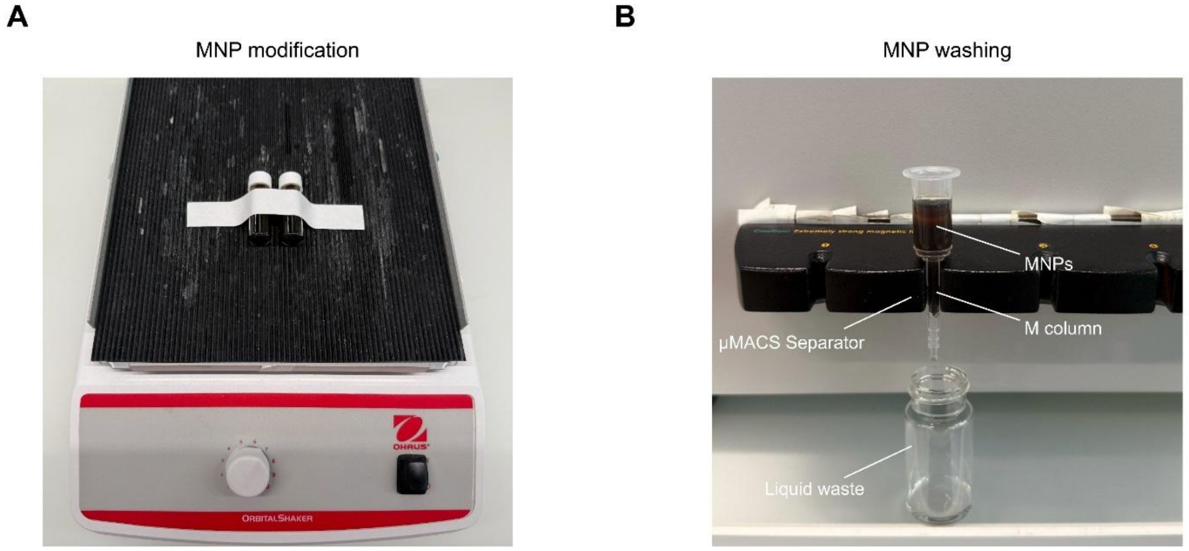

**Figure S15. Setup for MNP modification and washing. (A)** MNPs are being incubated with HaloTag ligands on an orbital shaker. **(B)** After modification, MNPs are being washed using an M column mounted on a μMACS Separator.
